# Low Dose Methotrexate has Divergent Effects on Cycling and Resting Human Hematopoietic Stem and Progenitor Cells

**DOI:** 10.1101/2025.02.19.639055

**Authors:** Maximilien Lora, Henri A. Ménard, Anastasia Nijnik, David Langlais, Marie Hudson, Inés Colmegna

## Abstract

Low dose methotrexate (LD-MTX) remains the gold standard in rheumatoid arthritis (RA) therapy. Multiple mechanisms on a variety of immune cells contribute to the anti-inflammatory effects of LD-MTX. Inflammatory signaling is deeply implicated in hematopoiesis by regulating hematopoietic stem and progenitor cell (HSPC) fate decisions; raising the question of whether HSPC are also modulated by LD-MTX. This is the first study to characterize the effects of LD-MTX on HSPC. CD34+ HSPC were isolated from healthy donors’ non-mobilized peripheral blood. Resting and/or cycling HSPCs were treated with LD-MTX [dose equivalent to that used in RA patients]. Flow cytometry was performed to assess HSPC viability, cell cycle, surface abundance of reduced folate carrier 1 (RFC1), proliferation, reactive oxygen species (ROS) levels, DNA double-strand breaks, p38 activation, and CD34+ subpopulations. HSPC clonogenicity was tested in colony-forming cell assays. Our results indicate that in cycling HSPC, membrane RFC1 is upregulated and, following LD-MTX treatment, they accumulate more intracellular MTX than resting HSPC. In cycling HSPC, LD-MTX inhibits HSPC expansion by promoting S-phase cell-cycle arrest, increases intracellular HSPC ROS levels and DNA damage, and reduces HSPC viability. Those effects involve the activation of the p38 MAPK pathway and are rescued by folinic acid. The effects of LD-MTX are more evident in CD34+CD38High progenitors. In non-cycling HSPC, LD-MTX also reduces the proliferative response while preserving their clonogenicity. In summary, HSPC uptake LD-MTX differentially according to their cycling state. In turn, LD-MTX results in reduced proliferation and the preservation of HSPC clonogenicity.

**Study Highlight Questions:** *What is the current knowledge on the topic?:*

- Low dose-methotrexate (LD-MTX) regulates the function of key cells involved in rheumatoid arthritis (RA) pathogenesis (e.g. T cells, macrophages, neutrophils, endothelial cells and fibroblast-like synoviocytes) and through the activation of multiple pathways contribute to the suppression of inflammation in RA.
- Inflammatory signaling impacts hematopoiesis by regulating hematopoietic stem and progenitor cells (i.e. HSPC, CD34^+^ cells).

*What question did this study address?:*

- Does LD-MTX modulate key functional properties of HSPC subpopulations (i.e. resting and/or cycling HSPCs)

*What does this study add to our knowledge?:*

- HSPC uptake LD-MTX differentially according to their cycling state.
- Cycling HSPC upregulate membrane RFC1 and uptake more MTX than resting HSPC with activation of the p38 MAPK pathway. This leads to inhibition of HSPC expansion, increased intracellular ROS levels and DNA damage, and reduced HSPC viability.
- LD-MTX preserves the clonogenicity of non-cycling HSPC.

*How might this change clinical pharmacology or translational science?:*

- We describe a novel mechanism of action of LD-MTX that extends its therapeutic effects from mature immune cells to the modulation of hematopoiesis.

## Introduction

Low-dose methotrexate (LD-MTX) is the mainstay initial treatment in rheumatoid arthritis (RA) and enhances the effect of biologic agents(1). LD-MTX promotes control of inflammation, prevents joint and organ damage, and reduces the risk of death among RA patients(2–4). However, despite its 40 years of clinical use there is an incomplete understanding of the cellular targets and molecular pathways underlying LD-MTX effects(5).

Hematopoietic stem and progenitor cells (i.e. HSPC, CD34^+^ cells) support hematopoiesis throughout life and generate all circulating immune cells. Inflammatory signaling is deeply implicated in hematopoiesis by regulating HSPC fate decisions(6). In mice with collagen induced arthritis, HSPC have broad downregulation of cell growth and proliferation genes expression. As a result, despite proliferative inflammatory signals, HSPC remain quiescent(7). Similar features are reported in RA patients, whose HSPC numbers and proliferative capacity are decreased and have a reduced response to growth factors(8–10). If such changes in the homeostatic control of hematopoiesis are relevant to RA immunopathogenesis, it raises the possibility that they could also be modulated by LD-MTX.

HSPC express the reduced folate carrier 1 (RFC1) transporter, encoded by the gene *SLC19A1*, required for MTX uptake(11). Before the widespread use of folic acid supplementation, the risk of pancytopenia in RA patients on LD-MTX was 1.4 to 2% (12). More recently, in the Cardiovascular Inflammation Reduction Study (CIRT), with the concomitant use of folic acid, the frequency of pancytopenia was reduced to 0.5% (13 cases). Furthermore, CIRT documented a modest protective effect of LD-MTX against thrombocytopenia(13). This suggests that LD-MTX can not only enter but also modulate HSPC. This paper explores that idea by, as a first step, characterizing in vitro the effects of LD-MTX on normal circulating HSPC.

## Methods

### HSPC isolation, culture, and LD-MTX treatment

This study was approved by the McGill University-Ethics Board (ERB protocol number 09-239 GEN) and written consent was obtained from all participants (healthy donors). Peripheral blood mononuclear cells (PBMC) from 40 ml of non-mobilized peripheral blood were isolated by gradient centrifugation using Lymphocyte Separation Medium (Mediatech Inc., Corning, NY, USA). CD34^+^ cells were labeled with the CD34 Microbead Kit human (Miltenyi Biotec Inc., Germany), magnetic–activated cell sorted with an AutoMACS (Miltenyi Biotec Inc., Germany), counted, and plated. HSPC numbers and viability were assessed by trypan blue exclusion on a hemocytometer. HSPC were seeded in 96-well flat-bottomed plates at 10 000 cells/well in 200µl of H3000 medium (StemSpan H3000; STEMCELL Technologies, Vancouver, BC, Canada) supplemented with penicillin/streptomycin. CC100 cytokine mix [human stem cell factor (SCF), human FMS-like tyrosine kinase 3 ligand (Flt3 ligand), human interleukin 3 and 6 (IL-3, IL-6); STEMCELL Technologies, Vancouver, BC, Canada] at the concentration suggested by the supplier (1X) was added to all cultures. ‘Quiescent’ HSPC were treated with LD-MTX (100nM; Methotrexate, Millipore Sigma, CA, USA) at day 0. In contrast, ‘proliferating’ HSPC were expanded for 2 days prior to adding LD-MTX. To antagonize the effect of MTX, 20 µM of folinic acid (FA, Leucovorin, Millipore Sigma, CA, USA) was added. CD34 depleted PBMC were cultured in complete RPMI1640 (Wisent Bioproducts, Saint-Jean-Baptiste, QC, Canada), supplemented with 10% FBS (Wisent Bioproducts, Saint-Jean-Baptiste, QC, Canada) and penicillin/streptomycin (Wisent Bioproducts, Saint-Jean-Baptiste, QC, Canada), and used for compensations and standards. HSPC and PBMCs were cultured in a humidified incubator at 37°C with 5% CO_2_.

### Flow cytometric analyses: RFC1, intracellular MTX, HSPC-proliferation, ROS, γH2AX, phospho-p38 and viability

To determine the levels of the MTX transporter reduced folate carrier 1 in HSPC (*HSPC-RFC1 levels)*, resting PBMCs were used as an internal control. Cells were treated with FcR blocking reagent (Miltenyi Biotech inc., Germany) and stained with surface antibodies [APC mouse anti-human CD34 (APC-CD34) and FITC mouse anti-human CD4 (FITC-CD4)] (BD Biosciences, Franklin Lakes, NJ, USA). After 30 min, cells were washed, fixed and permeabilized with Cytofix/Cytoperm Fixation/Permeabilization Solution Kit (BD Biosciences, Franklin Lakes, NJ, USA) following the manufacturer’s protocol. Rabbit anti-RFC1 (Alpha Diagnostic Intl Inc., San Antonio, TX, USA) was added for 1 hour. HSPC were washed and incubated for one hour at room temperature with the secondary antibody (PE goat anti-rabbit IgG, Cedarlane Laboratories Limited, Burlington, ON, Canada), washed and tested. Human RFC control/blocking peptide (Alpha Diagnostic Intl Inc., San Antonio, TX, USA) was used to optimize the concentrations of the primary and secondary anti-RFC1 antibodies. The flow cytometry gating strategy is shown in Figure S1.

To assess *MTX-HSPC uptake*, FITC-MTX (ThermoFisher Scientific, Waltham, MA, USA) at a final concentration of 1µM was added for 24 hours to HSPC cell cultures. HSPC were stained with APC-CD34 and the intensity of FITC-MTX in CD34^+^ cells was determined. The flow cytometry gating strategy is shown in Figure S2.

To assess *HSPC proliferation* we used the vital stain carboxyfluoroscein succinimidyl ester (CFSE, Millipore Sigma, CA, USA) fluorescent dye labeling as previously described(10). The expansion index (fold-expansion of the overall culture) was calculated with the FlowJo10 software. The gating strategy is shown in Figure S3. The HSPC *cell cycle assay* was assessed with Vibrant DyeCycle Violet stain (Molecular Probes Inc., Eugene, OR, USA), according to the manufacturer’s protocol. Intracellular *reactive oxygen species (ROS) in HSPC* were determined with 2’,7’-dichlorodihydrofluorescein diacetate (H_2_DCFDA, Molecular Probes, Eugene, OR, USA). HSPC were washed, resuspended in 10µM H_2_DCFDA in PBS, incubated for 20 minutes at 37°C in the dark and washed. HSPC were stained with APC-CD34 and BV605 mouse anti-human CD38 (BV605-CD38, BD Biosciences, Franklin Lakes, NJ, USA) followed by 7-AAD. The geometric mean fluorescence intensity (gMFI) of H_2_DCFDA on CD34^+^ and 7-AAD^-^ HSPC was recorded. The flow cytometry gating strategy is shown in Figure S4.

DNA double-strand breaks in *CD34 subpopulations were determined with γH2AX*. HSPC were stained with APC-CD34 and PE mouse anti-CD38 (BD Biosciences, Franklin Lakes, NJ, USA) for 30 min, washed and stained with 7-AAD for 20 min at room temperature. HSPC were fixed with 4% paraformaldehyde buffer for 10 min at room temperature, washed, permeabilized with Triton X-100 solution at 0.1% and incubated with AlexaFluor488 mouse anti-γH2AX (pS139, BD Biosciences, Franklin Lakes, NJ, USA) for 1 hour, washed and tested. The flow cytometry gating strategy is shown in Figure S5.

To assess *phospho-p38* HSPC were collected and fixed with BD Phosflow Fix buffer I and permeabilized with Perm Buffer III (BD Biosciences, NJ, USA) following manufacturer’s instructions. HSPC were incubated with APC mouse anti-human CD34 (APC-CD34), BV605-CD38, Pacific Blue mouse anti-phospho-p38 MAPK (pT180/pY182) (all from BD Biosciences, Franklin Lakes, NJ, USA). The Pacific Blue-p38 gMFI of HSPC was used to calculate fold increase levels (i.e., gMFI of HSPC treated with LD-MTX / gMFI of untreated HSPC). The flow cytometry gating strategy is shown in Figure S6.

All flow cytometric data were acquired with the BD LSR Fortessa (BD Biosciences) and analyzed with FlowJo software V10. *Viable* HSPC were defined as 7AAD negative. Only viable HSPC were analyzed in non-fixed datasets.

### Colony-forming cell (CFC) assay

Freshly isolated CD34^+^ cells were suspended in Methocult H4034 Optimum media (STEMCELL Technologies, Vancouver, BC, Canada) in duplicates and processed according to the manufacturer’s instructions. After 14 days of culture, the colonies of colony-forming unit-granulocyte macrophage (CFU-GM), burst-forming unit-erythroid (BFU-E), and CFU-granulocyte-erythrocyte-monocyte-megakaryocyte (CFU-GEMM) were counted with an inverted microscope.

### Statistical analyses

GraphPad Prism 10 software (GraphPad Software Inc., San Diego, CA, USA) was used for statistical analysis. Paired t-test was used to compare resting versus cycling HSPC and control versus MTX treated HSPC. Friedman test was used for the analysis of CD34 subpopulations. Two-way ANOVA was used to analyze all other datasets. Data are reported as mean ± standard deviation (SD). Significance was defined as *p < 0.05, **p < 0.01, or ***p < 0.001.

## Results

### In cycling HSPC, RFC1-mediated MTX-uptake promotes HSPC cycle arrest and death

HSPC self-renewal and multipotency capacities are central to maintain hematopoiesis(14). Self-renewal sustains the stem cell population, while through proliferation and differentiation multipotent progenitors are generated. Several factors within the bone marrow niche control HSPC’s dynamic. To assess the effect of MTX treatment in cycling HSPC, we used a combination of early- and late-acting recombinant human cytokines that support HSPC proliferation (i.e. CC100 cytokine cocktail: SCF, Flt3 ligand, IL-3, and IL-6). HSPC from non-mobilized peripheral blood are quiescent (i.e. G_0_ phase of the cell cycle), following culture in CC100 they restart cycling and proliferate for 2 days in culture. The dose of MTX (100nM) was informed by previous studies showing that this concentration is equivalent to that in human plasma following LD-MTX treatment(15). Further, we performed dose response experiments specifically using HSPC that confirmed the adequacy of the 100nM MTX dose (Figure S7). Cycling HSPC upregulate the MTX transporter reduced folate carrier 1 (RFC1) (Figure 1A) required for MTX uptake. This results in higher HSPC intracellular-MTX levels following LD-MTX treatment (Figure 1B). In addition, LD-MTX treatment of cycling HSPC halts their proliferation (Figure 1C) in S-phase (Figure 1D). This phenomenon is reversed by the addition of folinic acid (Figure 1C).

**Figure 1.**
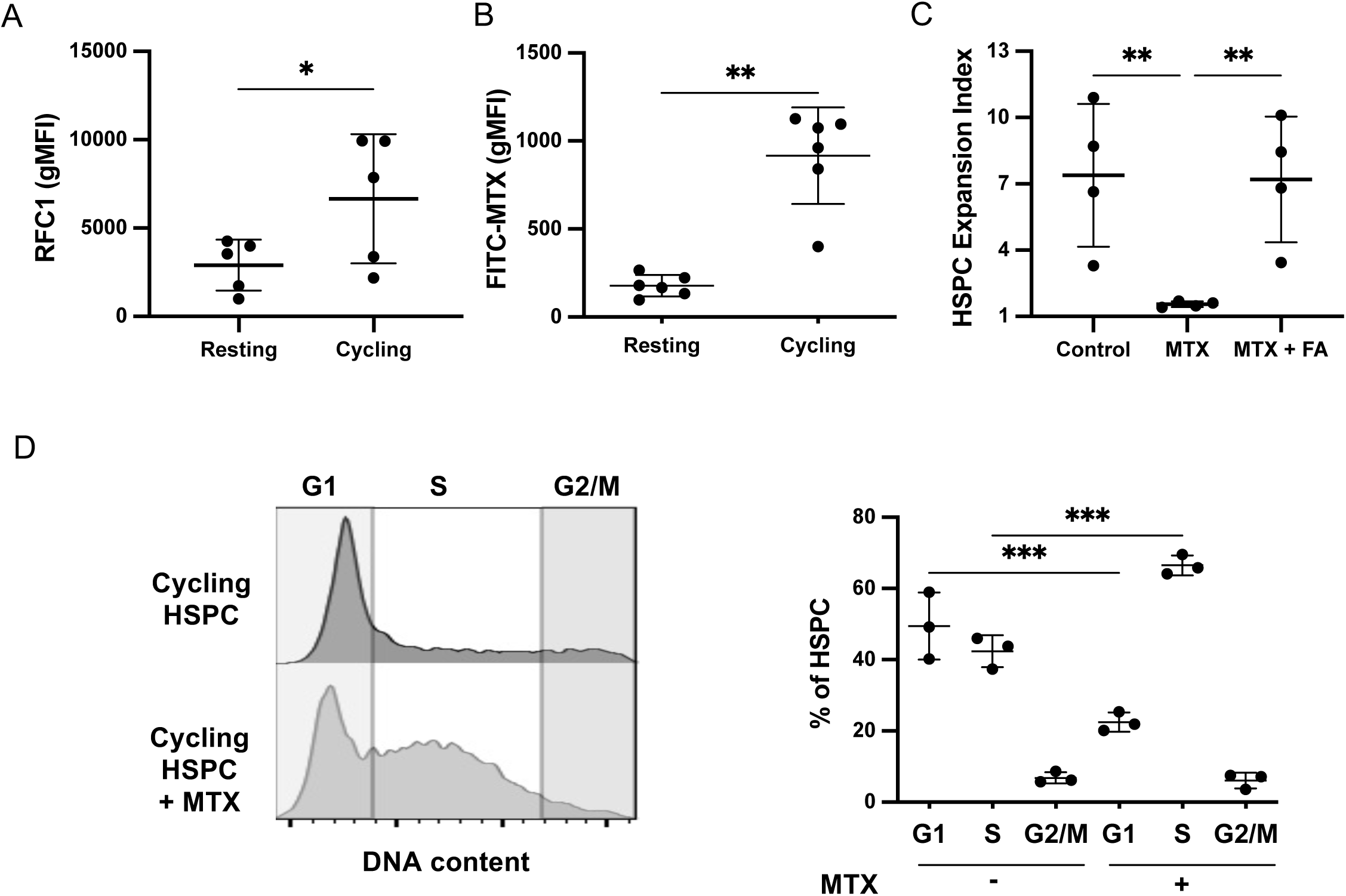
Low-dose methotrexate inhibits cycling HSPC expansion and promotes S-phase cycle arrest. (A) RFC1 in fixed permeabilized HSPC at day 0 (resting) and following 48hs of culture on StemSpan CC100 (cycling), n=5. (B) Intracellular MTX in resting and cycling HSPC following 24 hours treatment with 1 µM FITC-MTX, n=6. (C) Expansion index of cycling HSPC following LD-MTX treatment in the presence or absence of folinic acid, n=4. (D) Effect of LD-MTX on cycling HSPC cellular cycle (Vibrant DyeCycle Violet) representative example and quantification of frequency of HSPC in each phase of cell cycle, n=3. Mean ± SD. * p < 0.05; ** p < 0.01; *** p < 0.001.

### LD-MTX-induces apoptosis sensitization of cycling HSPC via p38 MAPK and JNK signaling

In lymphocytes and monocytes, LD-MTX not only interferes with cell-cycle progression but also increases apoptosis due to MTX-induced production of reactive oxygen species (ROS)(16). We tested if LD-MTX reduced the viability of cycling HSPC. Our results confirm that LD-MTX treatment promotes ROS production (Figure 2A), induces the formation of foci containing serine139-phosphorylated histone H2A.X (γH2A.X) (Figure 2B) and decreases the viability of cycling HSPC (Figure 2C). These findings support that MTX activates the ROS/DNA damage axis and induces apoptosis of cycling HSPC. Of interest, the effects on HSPC proliferation and viability uniquely follow LD-MTX treatment and do not occur when HSPC are treated with sulfasalazine, hydroxychloroquine, or ibuprofen (Figure S8).

**Figure 2.**
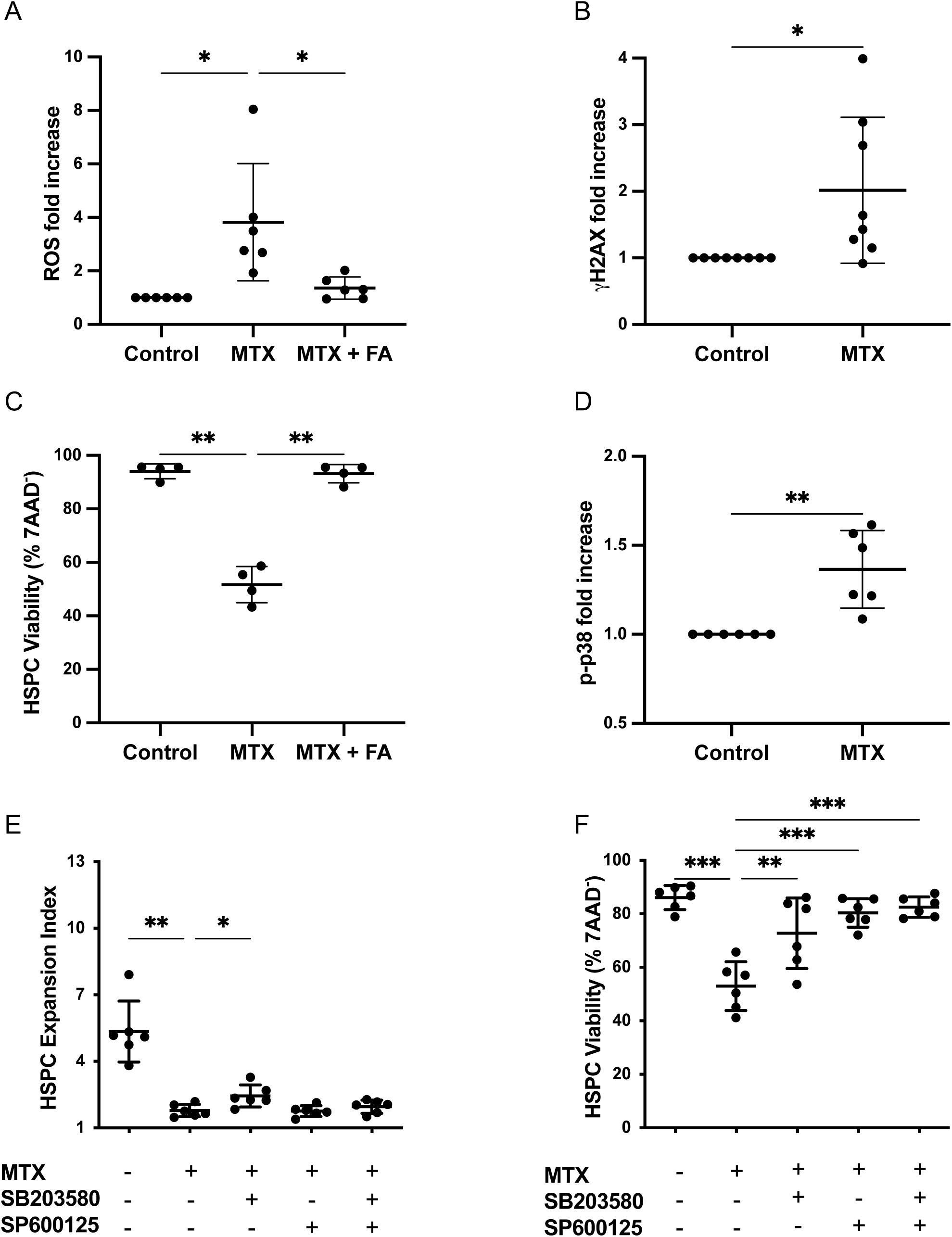
Low-dose methotrexate causes oxidative stress, DNA damage, p38 and JNK activation and reduced viability of cycling HSPC. All the following readouts were determined by flow cytometry in cycling HSPC: (A) ROS levels (H_2_DCFDA), n=6; (B) γH2AX, n=8; (C) viability [following 4 days of culture in StemSpan CC100 cocktail with MTX (100 nM) added 2 days after initiation of culture with or without FA (20 µM)], n=4; (D) Phospho-P38 [following 4 days of culture in CC100 with MTX (100 nM) added 2 days after initiation of culture with or without SB203580 (40 µM) and SP600125 (45µM)], n=6; (E) Expansion Index (CFSE), and (F) viability. Mean ± SD. * p< 0.05, ** p< 0.01, *** p< 0.001.

Mitogen-activated protein kinases (MAPKs) mediate cellular responses to various extracellular signals. Among them, p38 MAPK is activated by aberrant ROS levels in HSPC and can signal HSPC apoptosis, reducing long-term engraftment after transplantation (17,18). In contrast, inhibiting JNK signaling enhances the self-renewal of human HSPC(19). We tested if p38 and JNK signaling mediated the effects of LD-MTX treatment on HSPC. LD-MTX increases p38 phosphorylation in HSPC (Figure 2D). In the presence of a p38 inhibitor (SB203580, Adezmapimod, Calbiochem, San Diego, CA), HSPC expansion minimally improves and HSPC viability is restored (Figure 2E, F). In contrast, the use of a JNK inhibitor (SP600125, Calbiochem, San Diego, CA), does not affect HSPC proliferation but also improves HSPC viability (Figure 2E, F). These results support the involvement of MAPKs, in particular p38 and JNK activation in LD-MTX induced HSPC apoptosis.

### Stem and progenitor cells’ subpopulations differently respond to LD-MTX

CD38 is a type II membrane surface glycoprotein expressed on a variety of mature hematopoietic cells(20). CD38 expression is either low or absent on early HSPC populations and most primitive, pluripotent human hematopoietic stem cells are contained within the CD34^+^CD38^−^ fraction(21). Therefore, CD34^+^CD38^−^ cells long-term proliferation and blood cell production capacities exceed those of unfractionated CD34^+^ cells. We compared the effects of LD-MTX on HSPC subpopulations (i.e. CD38**^Low^** and CD38**^High^** gating strategy: Figure S2, S4, S5-S6). LD-MTX treatment increases ROS levels (Figure 3A), induces DNA damage (Figure 3B), activates p38 (Figure 3C), and reduces HSPC viability (Figure 3D) specifically in the CD34^+^CD38**^High^** subset where the uptake of FITC-MTX is highest (Figure 3E). This suggests an increased susceptibility of progenitors in contrast to stem hematopoietic subpopulations to the effect of LD-MTX.

**Figure 3.**
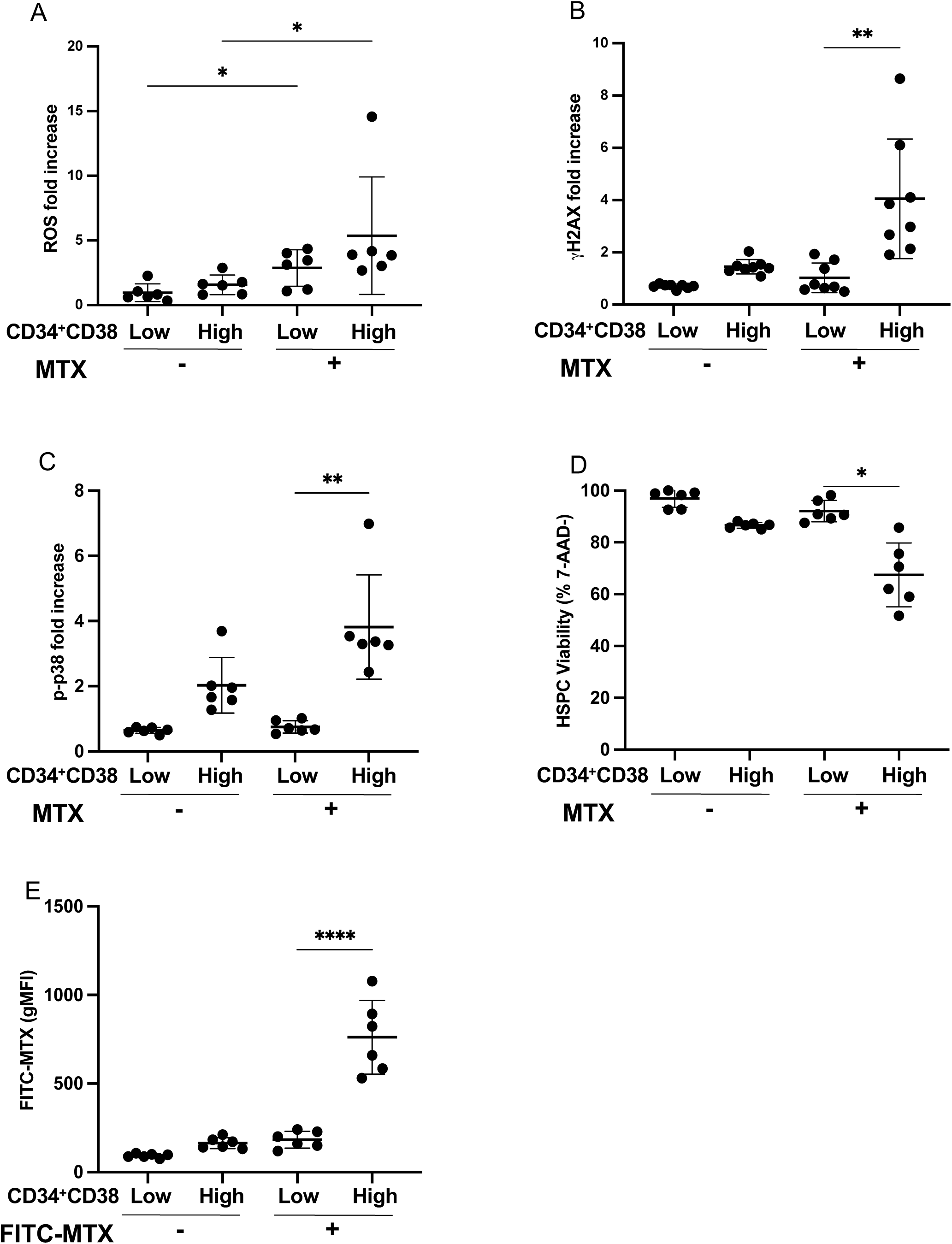
Low-dose methotrexate preferentially affects cycling CD38^High^ HSPC subpopulations. HSPC were expanded for 4 days with CC100 and MTX 100 nM was added 2 days after initiation of culture. The following readouts were determined by flow cytometry in CD34^+^CD38^Low^ vs CD34^+^CD38^High^ subpopulations: (A) ROS (H_2_DCFDA), n=6; (B) γH2AX levels, n=8; (C) Phospho-p38, n=6;(D) viability (CD34^+^/7AAD^-)^, n=6 and (E) FITC-MTX uptake, n=6. Mean ± SD. * p< 0.05, ** p< 0.01, **** p< 0.0001.

### LD-MTX treatment preserves the clonogenicity of quiescent HSPC

HSPC exist in both, cycling and G_0_ quiescent states(22). While cycling supports hematopoiesis, quiescence preserves the stem cell population. Therefore, quiescent HSPC (i.e., Day 0 or resting) have the highest clonogenic capacity(23) and following cycling (i.e., Day 4 of hematopoietic cytokine treatment) HSPC differentiate losing clonogenicity. To test if LD-MTX treatment affects HSPC clonogenic capacity, we stimulated HSPC proliferation in the presence or absence of LD-MTX. MTX treatment of resting HSPC, reduces the proliferative effect of cytokine treatment (Figure 4A), does not impact HSPC viability (Figure 4B), and preserves HSPC clonogenicity without skewing the ability to form specific types of colonies (Figure 4C-H). Altogether, these data indicate that cytokine-induced proliferation of resting HSPC is inhibited in the presence of MTX leading to clonogenic preservation.

**Figure 4.**
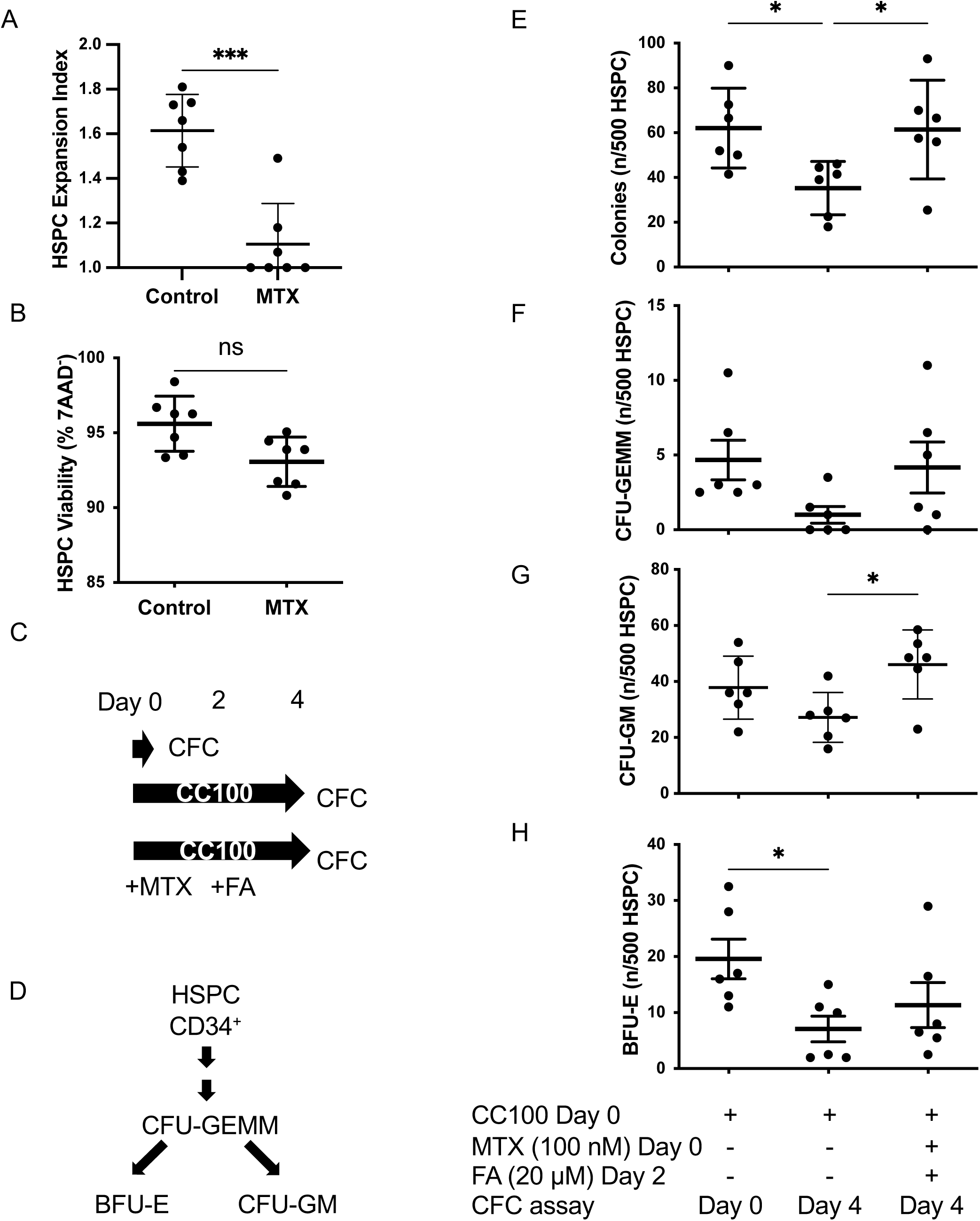
Low-dose methotrexate treatment preserves the clonogenicity of resting HSPC. Freshly isolated (resting) HSPC were treated or not with MTX (100 nM) and cultured for 2 days with StemSpan CC100 cocktail. HSPC (A) expansion index, n=7 and (B) viability, n=7 were assessed at day 2. The colony forming cell assay (CFC) was designed as in (C) to assess HSPC clonogenic potential (D). Total and specific types of colonies were counted: (E) total; (F) CFU-GEMM; (G) CFU-GM; and (H) BFU-E colonies per 500 HSPC. Mean ± SD. * p< 0.05; *** p< 0.001.

## Discussion

The success of low-dose MTX as an antirheumatic agent rests on its broad actions affecting a wide variety of pathogenic mechanisms resulting in anti-inflammatory effects. LD-MTX is up taken by several immune effectors (e.g., innate and adaptive immune cells, synovial fibroblasts, endothelial cells, bone and cartilage progenitors) particularly while they proliferate (i.e. cycling). The cycling activity of HSPCs is a result of extrinsic and intrinsic regulation(24). Infectious and inflammatory challenges promote HSPCs to proliferate and self-renew. This mechanism rapidly amplifies innate immune responses upon demand(25). On the other hand, proliferative stress is a driver of HSPC dysfunction. To test the effects of LD-MTX on cycling HSPC, we used recombinant human cytokines, an accepted approach to sustain ex-vivo HSPCs allowing for their functional assessment. Our results show that HSPC are equipped with RFC1 which is upregulated upon proliferation induction. As a result, HSPC uptake and likely polyglutamate MTX leading to a steady intracellular state(3). The intracellular levels of MTX in HSPC are a function of their activation state (i.e., low in resting and higher in cycling HSPC).

LD-MTX impacts key HSPC properties. In the presence of LD-MTX, cycling HSPC undergo S phase cycle arrest. This effect is reversed by folinic acid. Since quiescent HSPC have the highest self-renewal potential among all other blood cells(26), MTX-induced HSPC cycling arrest preserves their clonogenicity. This is particularly important since chronic inflammation and/or tonic levels of inflammatory signaling sustained over time and combined with aging negatively impact HSPC function (e.g., survival, proliferation, differentiation, self-renewal, and genomic integrity)(27). Moreover, accelerated inflammation-induced HSC cycling can enhance mutational occurrence. It is suggested that selective pressure imposed by chronic inflammation on the HSPC pool might drive genetic mutations and/or selection/expansion of inflammation-adapted mutant clones, leading to preleukemic states or leukemia(28). RA patients have an increased risk of all-cancer (20% relative excess risk) and hematological cancers compared to the general population(29). Disease activity, rather than RA treatment, seems to be the main driver of this risk(30). The fact that LD-MTX suppresses HSPC proliferation provides a direct mechanism that might reduce the HSPC mutagenesis risk.

LD-MTX enhances the production of reactive oxygen species (ROS), oxidative damage to DNA, activation of p38 and apoptosis sensitivity of cycling HSPC. ROS are toxic byproducts of aerobic metabolism that are harmful to living cells including HSPC, leading to DNA damage, senescence, or cell death. In rapidly dividing cell lines, MTX-induced ROS production mediates anti-inflammatory actions(31) and leads to γH2A.X phosphorylation, a marker of direct persistent double stranded DNA damage(32, 33). Several studies have shown that in HSPC, DNA damage and oxidative stress activate p38 which can induce HSPC apoptosis (17, 34–36). This prompted us to evaluate this pathway in HSPC following LD-MTX treatment. We showed that HSPC accumulate DNA breaks and exhibit high levels of intracellular ROS. The combination of these two features is key as accrued DNA damage itself is largely compatible with normal HSPC function if it does not result in massively elevated ROS(36). Given that JNK is crucial to regulate the expansion of human hematopoietic stem cells and that JNK inhibitors expand human stem cells with long-term repopulating capacity(19, 37), we assessed whether JNK inhibitors reversed the MTX-induced HSPC proliferative arrest. Our data suggest that the HSPC cycle arrest induced by LD-MTX involves p38 but not JNK activation. In contrast, both pathways are implicated in the reduced HSPC viability that follows LD-MTX treatment.

The first cell-surface marker used to enrich human HSPC was CD34, a ligand for L-selectin present in only 0.5–5% of hematopoietic cells in adult BM. Although almost all CD34^+^ cells have multi-potency or oligo-potency, this population is very heterogeneous. The surface marker CD38 allows enrichment of specific subpopulations. Most of CD34^+^ cells (90–99%) co-express CD38. However, the CD38 low to negative fraction contains cells that can give rise to multi-lineage colonies containing both lymphoid and myeloid cells(38). As such, CD34^+^CD38^-^ and not CD34^+^CD38^+^ are highly enriched for long-term culture-initiating cells(39). We assessed the effects of MTX in cycling CD38**^Low^** and CD38**^High^** HSPC populations. Hematopoietic stem cells (CD34^+^CD38**^Low^**) produce less ROS, have lower DNA damage, reduced p38 activation and higher viability than the more differentiated progenitors. This may relate to a differential response of both HSPC subtypes to cytokines-induced proliferation with reduced cycling in less differentiated HSPC and thus reduced MTX uptake in this subset.

This study has limitations. The number of circulating HSPC is small (i.e. ∼80 x10^3^) and we purposely did not expand them to avoid in vitro artefacts. For this reason, we increased the number of samples tested and not every dataset was generated with the same HSPC. We studied HSPC from healthy donors as the frequency of circulating HSPC in RA patients is even lower(10). We can’t rule out that other mechanisms are implicated in the response of RA-HSPC to LD-MTX. Finally, we did not assess potential effects of LD-MTX on adenosine signaling in HSPC. Adenosine regulates the generation of HSPC in the early embryo acting through A2b in endothelial cells(40). The role of adenosine in adult hematopoiesis is not yet defined.

To our knowledge, this is the first description of the effects of LD-MTX on HSPC. This work expands the targets of LD-MTX to hematopoietic stem and progenitor cells. Specifically, we demonstrate that HSPC uptake LD-MTX differentially according to their stemness, and that this results in key changes in two fundamental HSPC properties: self-renewal and differentiation. We propose mechanisms that mediate the effect of LD-MTX in HSPC, mainly oxidative stress and DNA damage, activating p38 and reducing HSPC viability. We show that a fundamental consequence of LD-MTX treatment of HSPC is the protection from proliferative exhaustion and the preservation of their clonogenicity. This may have significant consequences in terms of preventing the accumulation of genomic mutations in HSPC. Future studies should assess whether LD-MTX treatment reduces the oncogenic risk in RA by reducing the accumulation of HSPC genomic mutations.

## Acknowledgements

This work was supported by two grants from the Canadian Institutes of Health Research (CIHR) (CIHR PJT-173417, CIB-287233) and one from the Canadian Arthritis Network (11-02-PGP-03). Dr Langlais holds a Junior 2 salary award from the Fonds de recherches du Québec – Santé. We acknowledge Dr Linda Peltier’s assistance in the interpretation of the CFC assays.

## Author Contributions Statement

M.L., H.A.M., A.N., D.L., M.H. and I.C wrote the manuscript; M.L., H.A.M. and I.C. designed the research; M.L. and I.C. performed the research; M.L., A.N., D.L., M.H. and I.C analyzed the data; I.C. and M.H. contributed patient samples.

## Conflicts of interest

Authors have no conflict of interest to declare.

## Funding information

This work was supported by grants from the Canadian Arthritis Network (11-02-PGP-03), the Canadian Institutes of Health Research (CIHR) (CIHR PJT-173417, CIB-287233) and the Fonds de recherche du Québec – Santé (FRQS).

---

**Figure S1.**
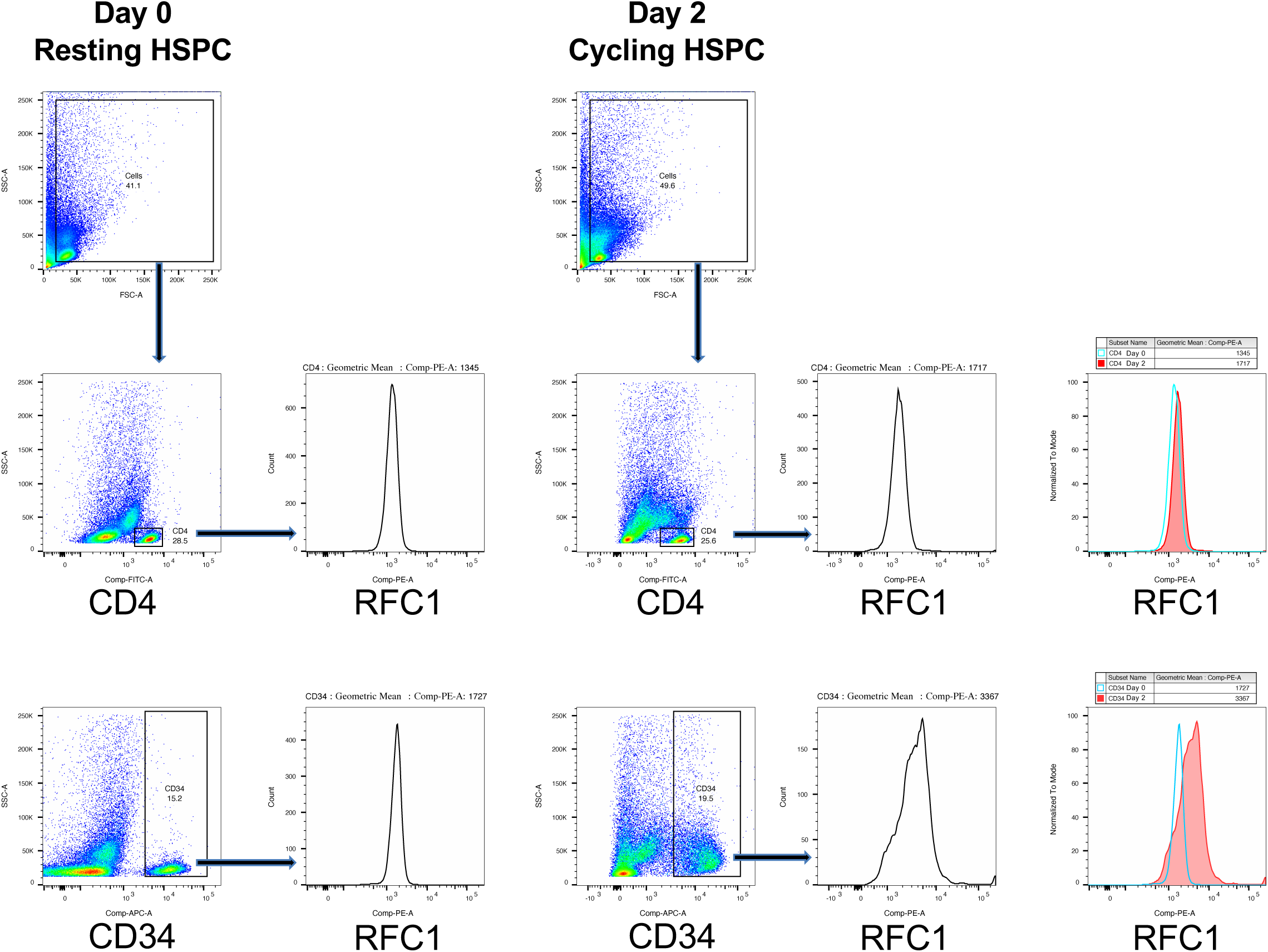
RFC1 quantification by flow cytometry (gating strategy). HSPC were cultured for 2 days with CC100 cocktail. To determine the levels of the RFC1 in HSPC, resting PBMCs (100 000 cells) were used as an internal control. Cells were treated with FcR blocking reagent and stained with surface antibodies [APC mouse anti-human CD34 (APC-CD34) and FITC mouse anti-human CD4 (FITC-CD4)]. After 30 min, cells were washed, fixed and permeabilized with Cytofix/Cytoperm Fixation/Permeabilization Solution Kit following the manufacturer’s protocol. Rabbit anti-RFC1 was added for 1 hour. HSPC were washed and incubated for one hour at room temperature with the secondary antibody (PE goat anti-rabbit IgG), washed again and analyzed by flow cytometry.

**Figure S2.**
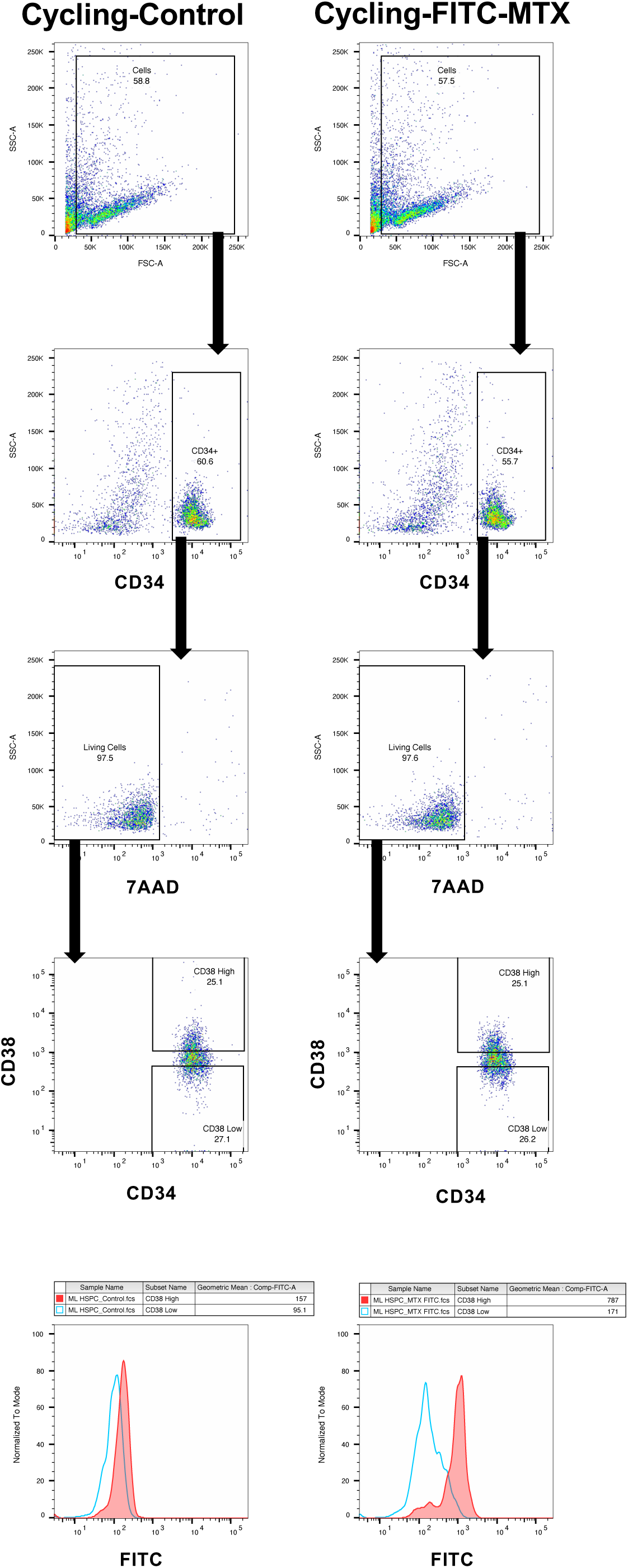
FITC-MTX quantification by flow cytometry (gating strategy). HSPC were cultured for 24 hours with FITC-MTX (1 µM) starting at day 0 for Resting and day 2 for Cycling samples. FITC-MTX at 100 nM was also tried to match the MTX conditions in all our experiments but the FITC fluorescence was barely detectable.

**Figure S3.**
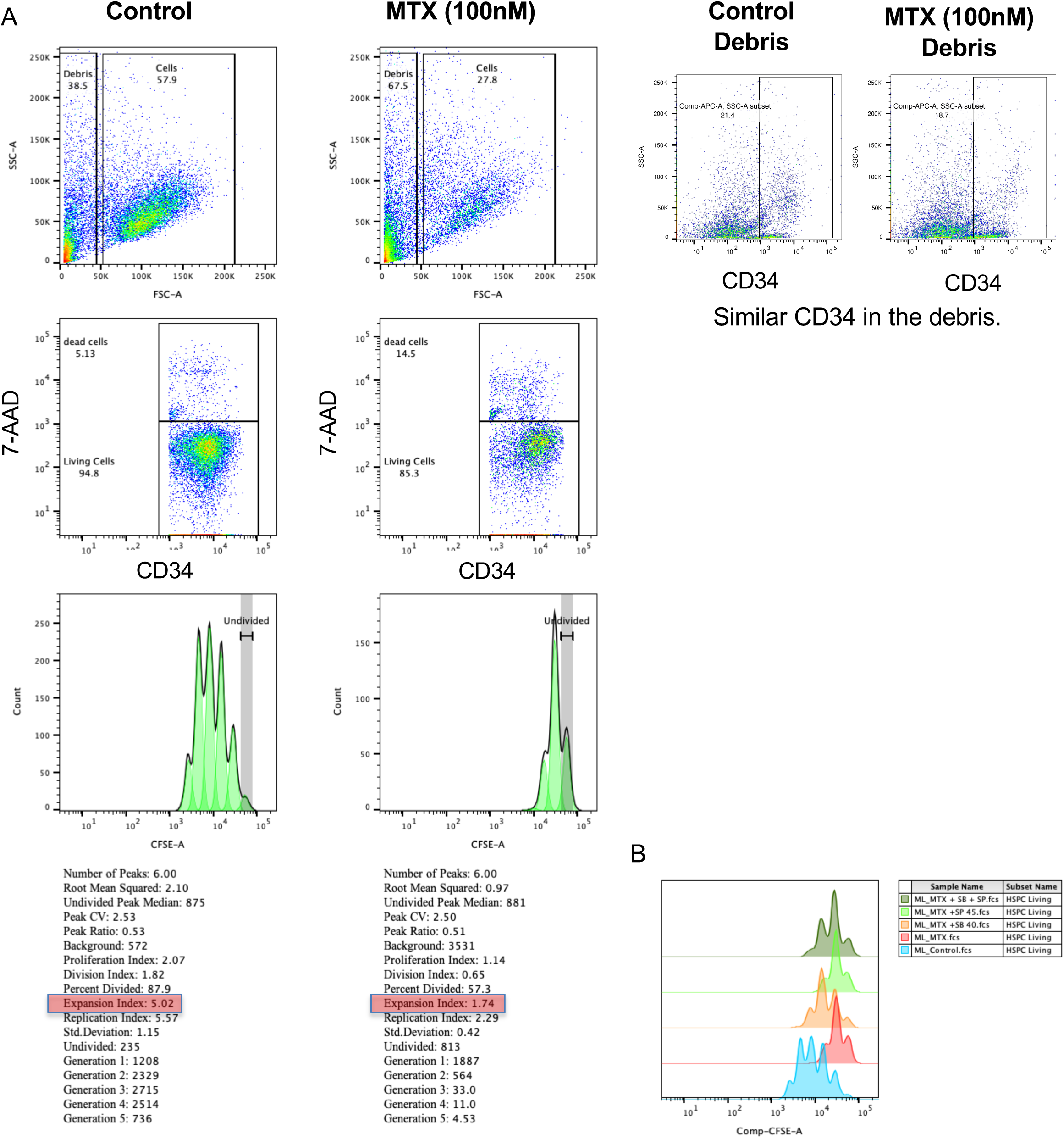
Proliferation (CFSE staining) of living HSPC in the presence of low-dose methotrexate (gating strategy). HSPC were cultured for 4 days with CC100 cocktail in the presence or absence of MTX 100 nM from day 2. (A) Expansion index was calculated in CFSE stained HSPC with FlowJo 10. (B) Representative example of the effects of different treatments on HSPC proliferation. Treating the HSPC with MTX in the presence of SB203580 40 µM (SB 40) or SP600125 45 µM (SP 45) and in combination (SB + SP).

**Figure S4.**
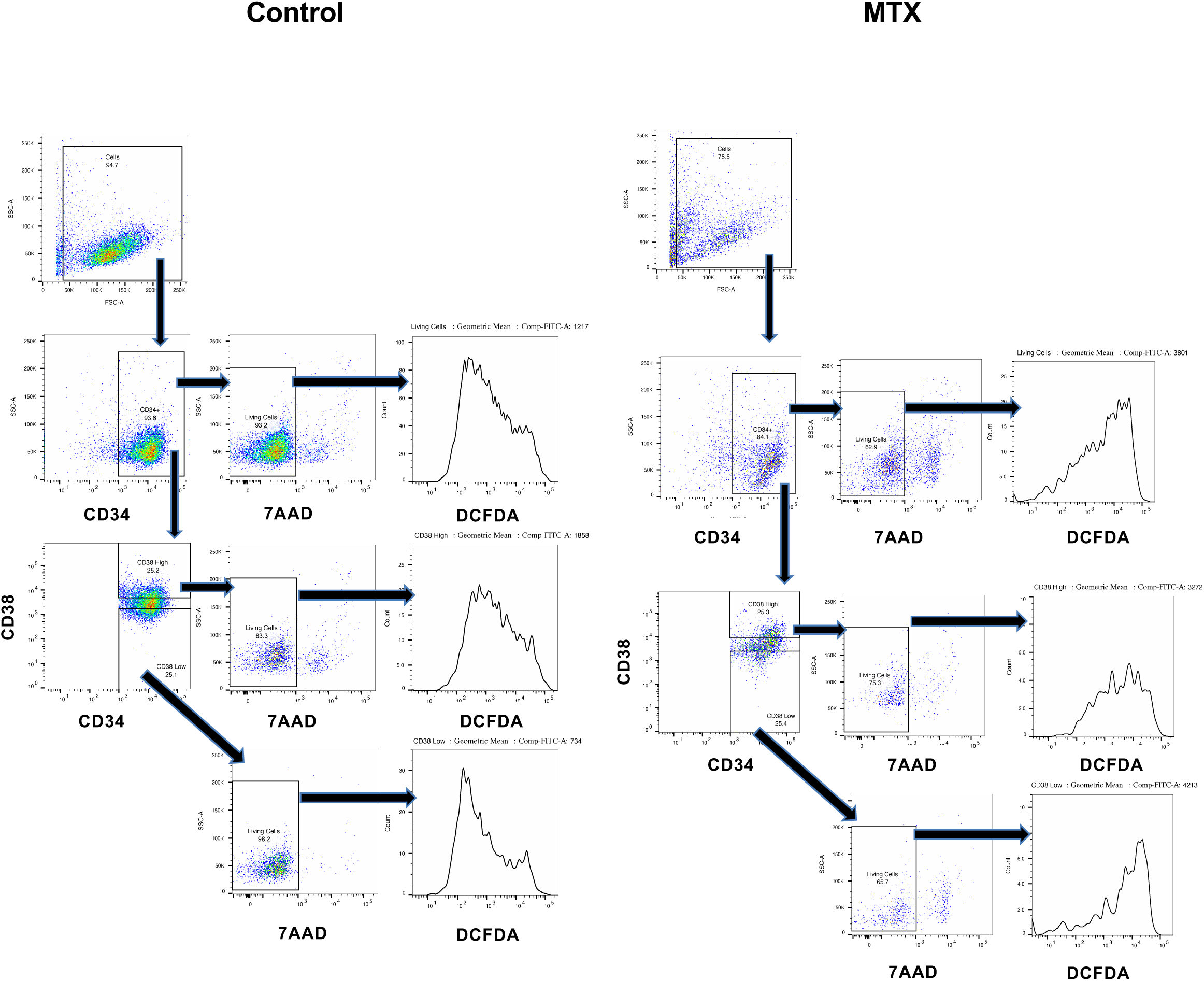
ROS quantification with H_2_DCFDA by flow cytometry (gating strategy). The assessment of intracellular *reactive oxygen species (ROS) in HSPC* was determined using 2’,7’-dichlorodihydrofluorescein diacetate (H_2_DCFDA, Molecular Probes, Eugene, OR, USA). HSPC were cultured for 4 days with CC100 cocktail with MTX (100 nM) added at the second day of culture. On day four, the HSPC were washed, resuspended in 10µM H_2_DCFDA in PBS, incubated for 20 minutes at 37°C in the dark and washed. HSPC were stained with APC-CD34 and BV605 mouse anti-human CD38 (BV605-CD38, BD Biosciences, Franklin Lakes, NJ, USA) followed by 7-AAD staining. The gMFI of DCFDA in CD34^+^CD38^low^ and CD34^+^CD38^high^ was calculated and expressed as fold increase compared to living CD34^+^ HSPC from the Control.

**Figure S5.**
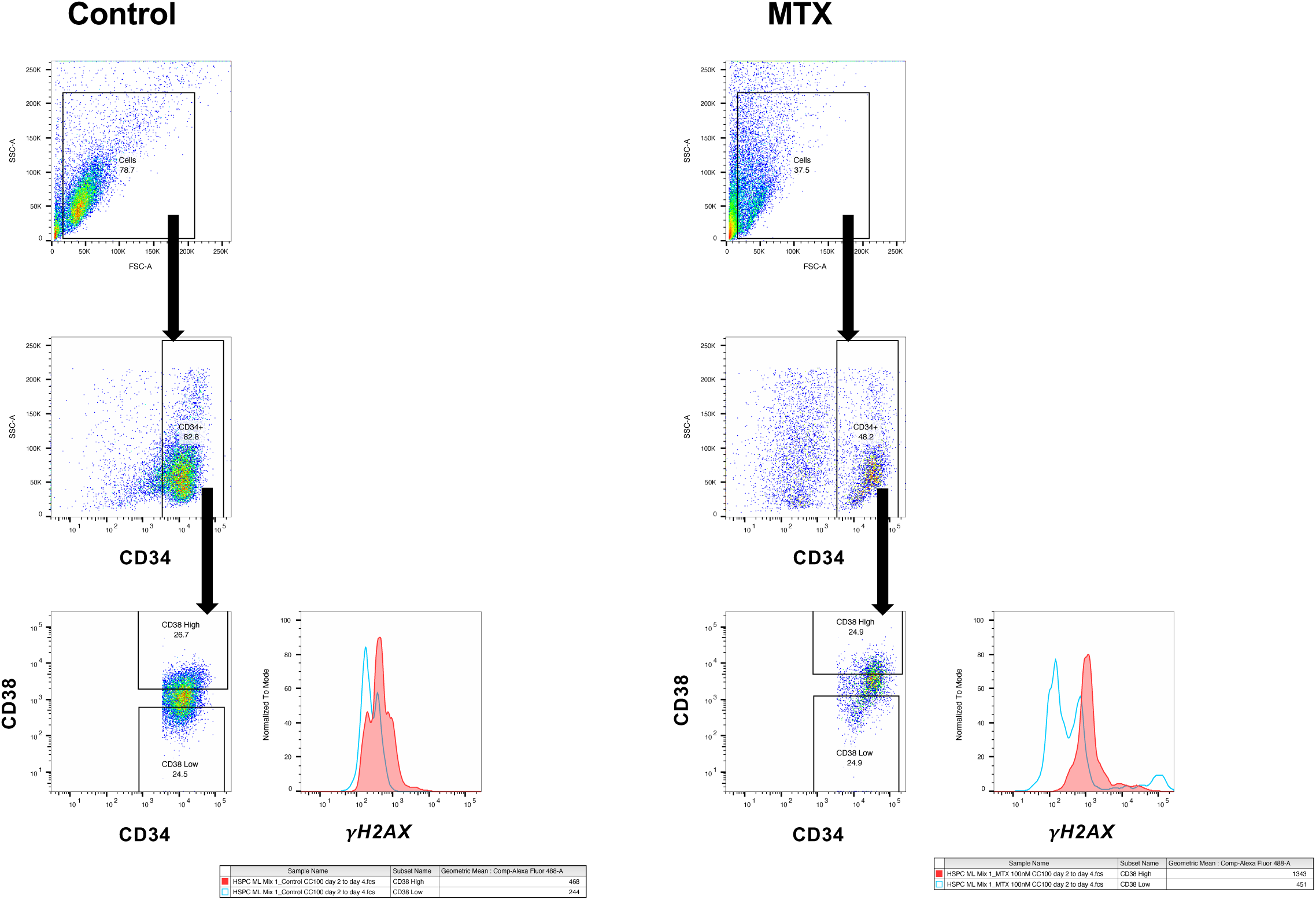
*γ*H2AX quantification by flow cytometry (gating strategy). *γH2AX* was used to determine DNA double-strand breaks in *CD34 subpopulations*. HSPC were cultured for 4 days with CC100 cocktail with MTX (100 nM) added at the second day of culture. On day four, HSPC were stained with cell surface markers APC-CD34 and PE-CD38 (BD Biosciences, Franklin Lakes, NJ, USA) for 30 min at 4°C and washed. HSPC were fixed with 4% paraformaldehyde fixation buffer for 10 min at room temperature, washed, permeabilized with Triton X-100 solution at 0.1% and incubated with AlexaFluor488 mouse anti-γH2AX (pS139, BD Biosciences, Franklin Lakes, NJ, USA) for 1 hour, washed again and analyzed by flow cytometry.

**Figure S6.**
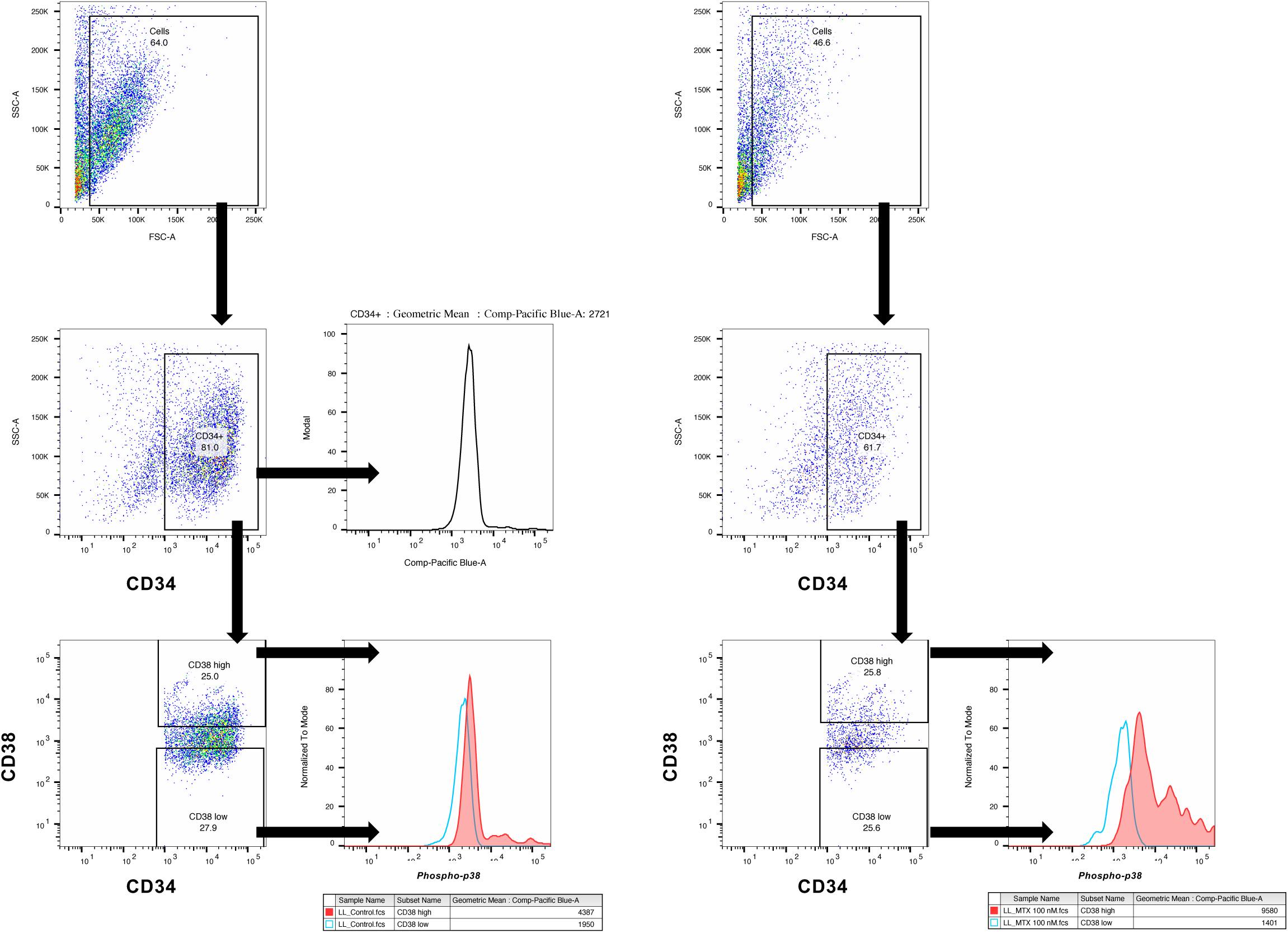
*Phospho-P38* quantification by flow cytometry (gating strategy). HSPC were cultured for 4 days with CC100 cocktail with MTX (100 nM) added at the second day of culture. To assess *phospho-p38,* HSPC were collected and fixed with BD Phosflow Fix buffer I and permeabilized with Perm Buffer III following manufacturer’s instructions. HSPC were incubated with APC-CD34, BV605-CD38, Pacific Blue mouse anti-phospho-p38 MAPK (pT180/pY182). The Pacific Blue-p38 gMFI of HSPC was used to calculate fold increase levels (i.e., gMFI of HSPC treated with LD-MTX / gMFI of untreated HSPC).

**Figure S7.**
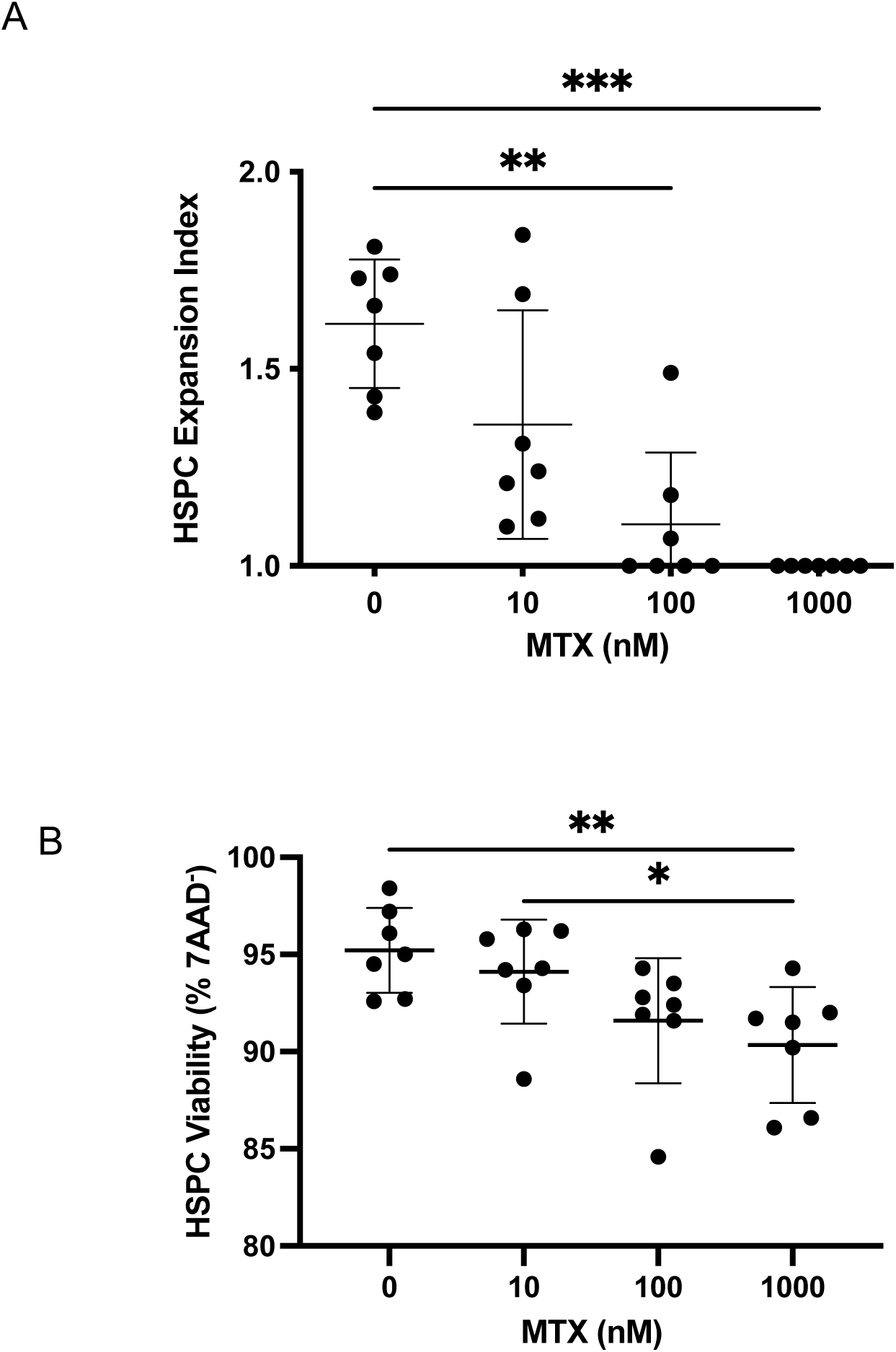
Dose response of low-dose methotrexate on HSPC proliferation and viability. HSPC were cultured for 2 days with CC100 and MTX at different concentrations added at the start of the culture. (A) HSPC expansion index was measured by CFSE dilution, and (B) viability was assessed by 7AAD. n=7. Mean ± SD. ** p< 0.01, *** p< 0.001

**Figure S8.**
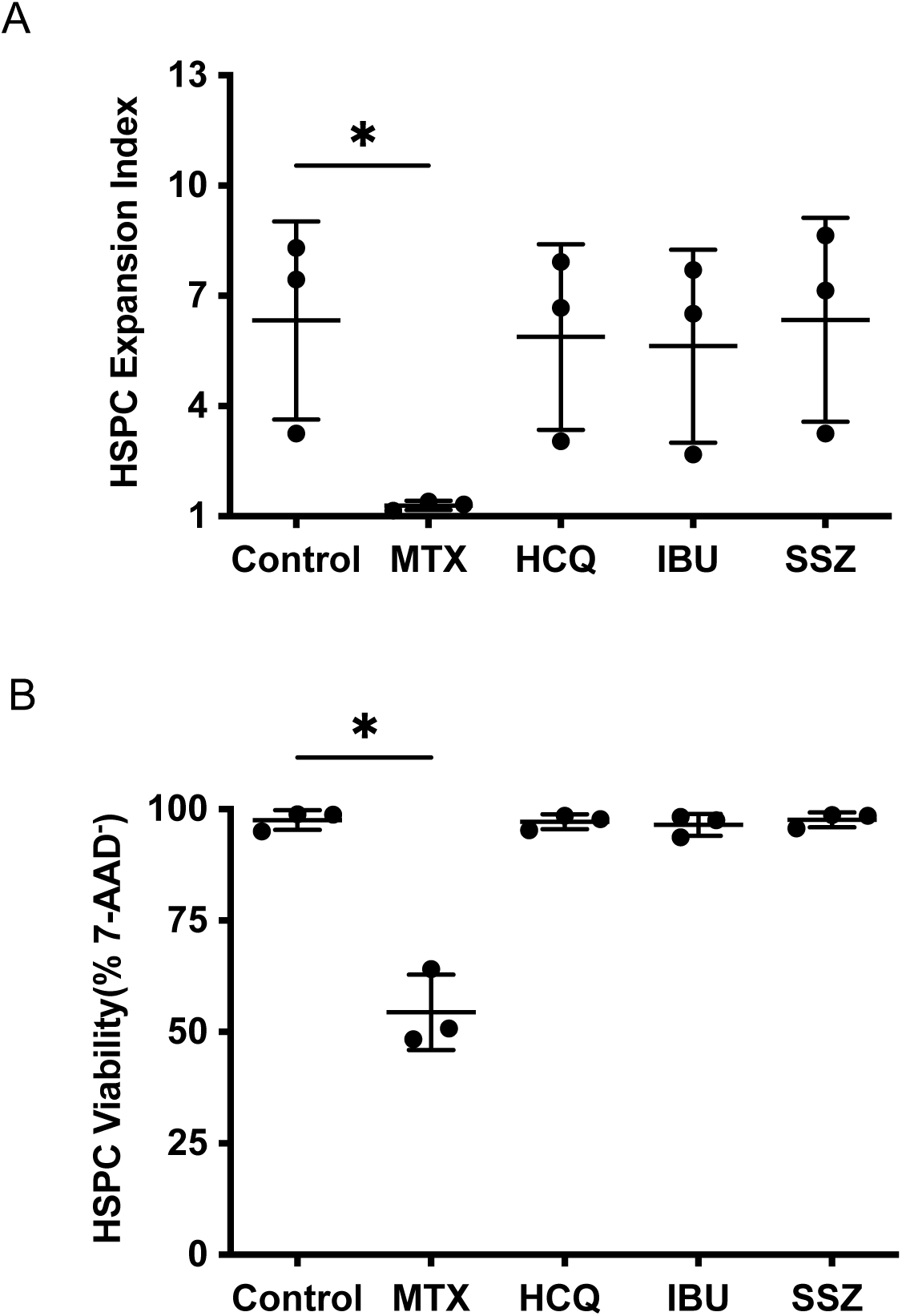
Low-dose methotrexate but not hydroxychloroquine, sulfasalazine or ibuprofen inhibits HSPC proliferation and reduces HSPC viability. HSPC were cultured for 4 days with CC100 cocktail in the presence of MTX 100 nM, HCQ 1 µM, IBU 100 µM and SSZ 10 µM. (A) HSPC expansion index was measured by CFSE staining, n=3, and (C) HSPC viability by 7-AAD incorporation, n=3. Mean ± SD. * p< 0.05

## Notes

### Competing Interest Statement

The authors have declared no competing interest.

